# A combination of physicochemical tropism and affinity moiety targeting of lipid nanoparticles enhances organ targeting

**DOI:** 10.1101/2023.11.21.568061

**Authors:** Marco E. Zamora, Serena Omo-Lamai, Manthan N. Patel, Jichuan Wu, Evguenia Arguiri, Vladmir Muzykantov, Jacob Myerson, Oscar Marcos-Contreras, Jacob S. Brenner

**Affiliations:** Drexel University; University of Pennsylvania

## Abstract

Two camps have emerged in the targeting of nanoparticles to specific organs and cell types: affinity moiety targeting, which conjugates nanoparticles to antibodies or similar molecules that bind to known surface markers on cells; and physicochemical tropism, which achieves specific organ uptake based on the nanoparticle’s physical or chemical features (e.g., binding to endogenous proteins). Because these camps are largely non-overlapping, the two targeting approaches have not been directly compared or combined. Here we do both, using intravenous (IV) lipid nanoparticles (LNPs) whose original design goal was targeting to the lungs’ endothelial cells. For an affinity moiety, we utilized PECAM antibodies, and for physicochemical tropism, we used cationic lipids, both having been heavily studied for lung targeting. Surprisingly, the two methods yield nearly identical levels of lung uptake. However, aPECAM LNPs display much greater specificity for endothelial cells. Intriguingly, LNPs that possess both targeting methods had >2-fold higher lung uptake than either method alone. The combined-targeting LNPs also achieved greater uptake in already inflamed lungs, and greater uptake in alveolar epithelial cells. To understand how the macro-scale route of delivery affects organ targeting, we compared IV injection vs. intra-arterial (IA) injection into the carotid artery. We found that IA combined-targeting LNPs achieve 35% of the injected dose per gram (%ID/g) in the brain, a level superior to any other reported targeting method. Thus, combining affinity moiety targeting and physicochemical tropism provides benefits that neither targeting method achieves alone.

**Graphical Abstract:** 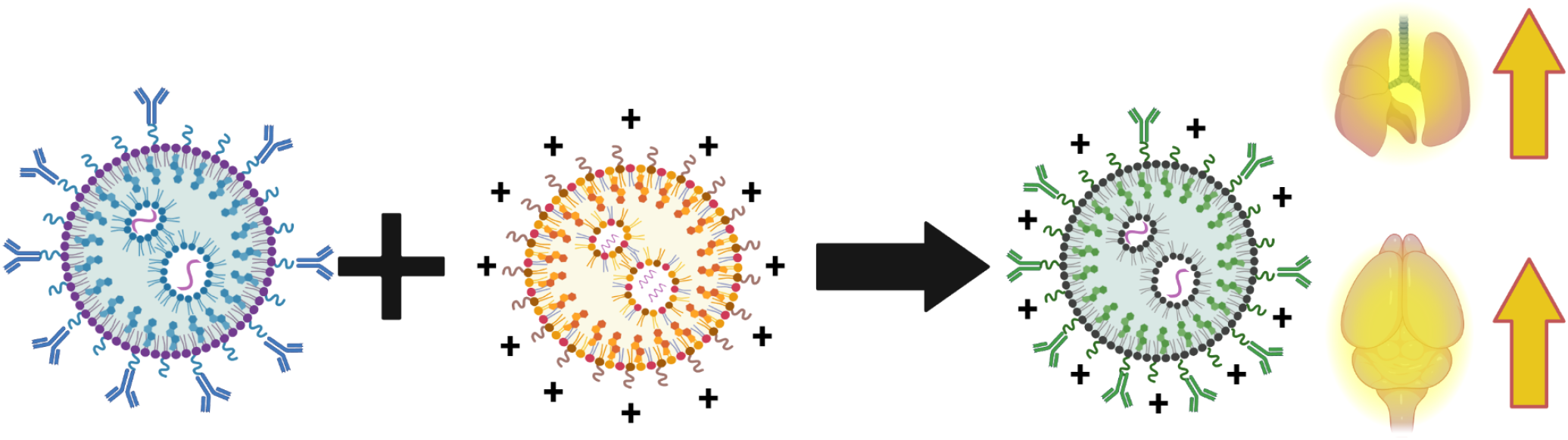

## Introduction

Nanoparticles have long held the promise of selectively targeting specific cell types and organs, though that goal has remained largely elusive for organs other than the liver, the natural clearance organ for intravenous nanoscale materials. In working to achieve the long-sought targeting goal in non-liver organs, two strategies have predominated, termed here “affinity moiety targeting” and “physicochemical tropism.”

“Affinity moiety targeting” was developed first, and employs “affinity moieties’’ which bind specifically to chosen epitopes on target cells; e.g., conjugating nanoparticles to monoclonal antibodies that have affinity for the target cell. This was first achieved by Torchilin and others, who showed the benefits of conjugating liposomes to monoclonal antibodies^1–6^. Numerous other affinity moieties have been used besides monoclonal antibodies, such as small molecules that bind to receptors on target cells, aptamers, fragments of antibodies (Fab, F(ab’)_2_, and many more ^7–16^. At least 10 nanoparticles targeted with affinity moieties have reached clinical studies, but none have yet been approved ^14,17,18^.

“Physicochemical tropism” selects nanoparticles for tropism to a particular organ based on the nanoparticle’s physical properties (size, shape, rigidity, charge) or chemical features (binding to specific endogenous proteins because of the nanoparticles’ chemical makeup). In general, physicochemical tropism is not easy to design from first principles, and therefore commonly it is achieved by screening large numbers of nanoparticles with slightly varying formulations in order to find one with tropism for a particular cell or organ. This screening-based approach is exemplified by lipid nanoparticle (LNP) screens, especially Anderson’s studies of the early 2010s^19–22^. Such screening, or optimized searches through LNP formulation space, have produced numerous LNP formulations with physicochemical tropism for a particular organ or cell type, ranging from the lungs to spleen, and achieving cell-type-specificity to macrophages and T-cells^23–27^. The first FDA-approved RNA-loaded lipid nanoparticle (LNP), patisiran, approved in 2018, has physicochemical tropism to the plasma protein ApoE, which selectively shuttles the LNPs to hepatocytes^28^.

Much has been written comparing affinity moiety targeting and physicochemical tropism^19–27,29–39^. Advantages cited for affinity moieties include a known mechanism of action, which enables rational engineering. Physicochemical targeting usually does not have a known mechanism of action (ApoE binding is a rare exception), and therefore rational engineering of a lead formulation is difficult. However, physicochemical targeting has a large advantage for industrialization, in that affinity moieties such as antibodies are not easy to conjugate to nanoparticles and then purify at scale^40,41^. Physicochemical targeting, for LNPs at least, usually just involves changing the lipid formulation, which is facile at industrial scale^42^. Thus, each approach has relative advantages and disadvantages. These two approaches have rarely been tested head-to-head, especially for LNPs. This lack of head-to-head testing might be because in general these two approaches are performed in separate labs, falling into separate “camps.”

Here, we directly compare cell and organ delivery efficiency and specificity when delivering to a single organ, the lung, using targeting to PECAM for affinity targeting vs. cationic lipids for physicochemical targeting. We do this for LNPs, since they are the leading nanoparticle format currently under development in industry, with tens of billions invested in the last few years^43,44^. We chose to target the lungs because many formulations with both affinity and physicochemical targeting have been created with the goal of ameliorating alveolar diseases, especially acute respiratory distress syndrome (ARDS), the alveolar inflammatory disease that was the primary cause of death during the COVID-19 pandemic, and still causes >75,000 deaths each year from non-COVID types of ARDS^45–50^.

To target LNPs to alveolar vasculature using affinity moieties, we chose LNPs conjugated to antibodies against PECAM-1. Compared to untargeted LNPs, PECAM targeting has been shown to increase mRNA delivery and protein expression in the lung by ∼200-fold and 25-fold respectively^31^. PECAM has been compared to other affinity moieties, and none perform markedly better with respect to lung uptake. For physicochemical tropism to the lungs, we chose LNPs that incorporate a cationic lipid. Numerous LNPs have been formulated that have physicochemical tropism to the lungs, with the commonality that nearly all possess lipids that bear a positive charge within some biologically-relevant pH range. Some of these are ionizable lipids with a tertiary amine, while others are permanently cationic lipids with a quarternary amine^23,25,51^. Here we chose the most cited such LNP formulation, which includes DOTAP, a permanently cationic lipid with a quaternary amine ^23–25,52^. DOTAP LNPs have been shown to drive expression of mRNA-luciferase primarily in the lungs^23–25,52^.

In comparing aPECAM and DOTAP LNPs, we find that they have nearly identical localization to the lungs, despite dramatically different targeting mechanisms. We then show that combining PECAM antibodies and DOTAP into a single LNP produces a synergistic benefit, achieving the highest reported delivery of an LNP to the lungs, and suggesting that bringing together the two approaches can produce higher uptake than either alone.

## Methods

### Materials

#### Lipids used for LNP formulations

DOTAP (1,2-dioleoyl-3-trimethylammonium-propane (chloride salt)), DOPE

(1,2-dioleoyl-*sn*-glycero-3-phosphoethanolamine), cholesterol, DMG-PEG 2000

(1,2-dimyristoyl-rac-glycero-3-methoxypolyethylene glycol-2000), DPPC (dipalmitoyl phosphatidylcholine), 18:1 PE TopFluor AF 594

(1,2-dioleoyl-sn-glycero-3-phosphoethanolamine-N-(TopFluor® AF594) (ammonium salt)), 18:0 PE-DTPA

(1,2-distearoyl-sn-glycero-3-phosphoethanolamine-N-diethylenetriaminepentaacetic acid (ammonium salt)), and DSPE-PEG2000-azide

(1,2-distearoyl-sn-glycero-3-phosphoethanolamine-N-[azido(polyethylene glycol)-2000] (ammonium salt)) were purchased from Avanti Polar Lipids. Ionizable lipid cKK-E12 was purchased from Echelon Biosciences.

#### Radiolabel

Indium-111 Chloride (In-111) was purchased from BWXT Medical.

#### Flow Cytometry surface marker antibodies

All antibodies described in **Supp Table 2** were purchased from Biolegend

### Nanoparticle production and protein modification

#### Production of fluorescent and radiolabeled LNP formulations

LNPs were formulated using the microfluidic mixing method. An organic phase containing a mixture of lipids dissolved in ethanol at a designated molar ratio (**Fig 1C and Supp Table 1**) was mixed with an aqueous phase (50 mM citrate buffer, pH 4) containing Luciferase mRNA (TriLink) at a flow rate ratio of 1:3 and at a total lipid/mRNA weight ratio of 40:1 in a microfluidic mixing device (NanoAssemblr Ignite, Precision Nanosystems). LNPs were dialysed against 1× PBS in a 10 kDa molecular weight cut-off cassette for 2 h, sterilized through a 0.22 μm filter and stored at 4 °C.

**Figure 1:**
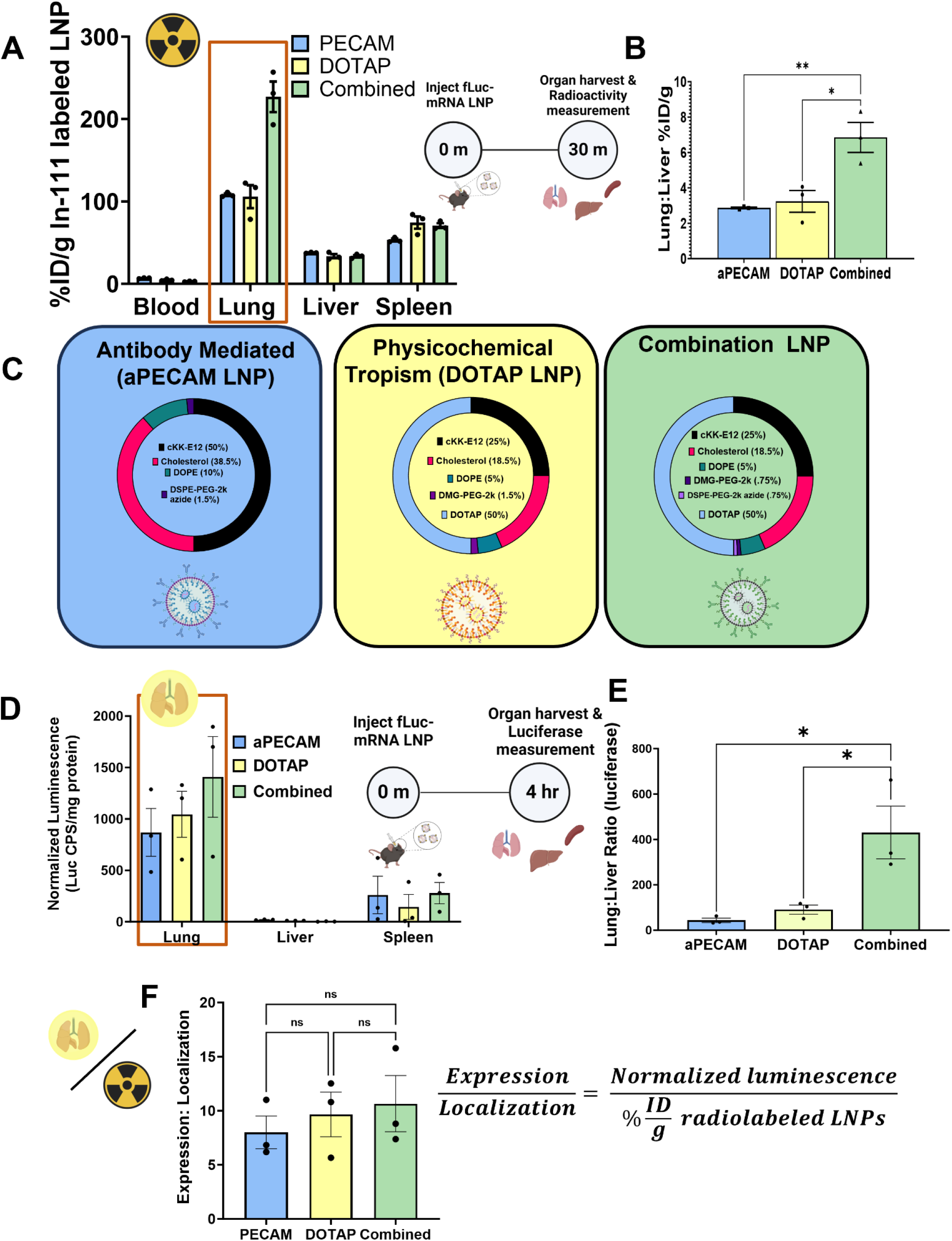
LNPs formulated with cationic lipids and targeted to PECAM show improved lung localization and expression. **A.** Biodistributions of aPECAM, DOTAP, and Combined LNPs 30 minutes after LNP injection. Combined LNPs localize to the lungs >2-fold more than aPECAM or DOTAP LNPs. **B.** Lung-to-liver ratios obtained from A calculated by dividing the %ID/g of tissue in the lung by that in the liver. Combined LNPs have a >2-fold higher lung-to-liver ratio compared to aPECAM or DOTAP LNPs. **C.** Formulations and molar percentages of aPECAM, DOTAP, and Combined LNPs. **D.** Luciferase expression of aPECAM, DOTAP, and Combined LNPs in the lung, liver, and spleen 4 hours after LNP injection. **E.** Lung-to-liver ratios calculated from D by dividing the normalized luciferase expression values in the lung by that in the liver. The lung-to-liver ratio of combined LNPs is about 10-fold and 5-fold higher than both aPECAM and DOTAP LNPs respectively. **F.** Ratios of lung expression to localization, calculated by dividing the normalized luciferase expression values from D to the %ID/g of tissue values from A for each LNP formulation. The expression to localization ratios are not significantly different between formulations indicating a good correlation between LNP localization and luciferase mRNA expression. **Statistics:** n=3 and data shown represents mean ± SEM. Comparisons between groups were made using 1-way ANOVA with Tukey’s post-hoc test. *=p<0.05, ***=p<0.001, ****=p<0.0001.

#### Nanoparticle Characterization

Dynamic light scattering measurements of hydrodynamic nanoparticle size, distribution, polydispersity index, and zeta potential were made using a Zetasizer Pro ZS (Malvern Panalytical) (**Supp Fig 1**). LNP RNA encapsulation efficiencies and concentrations were measured using a Quant-iT RiboGreen RNA assay (Invitrogen)

#### Antibody conjugation

Antibodies were conjugated to aPECAM and Combined LNPs by DBCO-Azide copper-free click chemistry. DBCO was added to the antibodies in 5 x molar excess and incubated at room temperature for 30 minutes, with rotation. To enable assessment of binding efficiency using fluorescence, the antibodies were fluorescently labeled using AF 647 fluorophore by incubating 10% of total antibody volume for 30 minutes at room temperature in the dark. The antibodies were then purified using Amicon centrifugal filters, and centrifuged at 3200 x g for 20 minutes to achieve a total flowthrough 10x antibody volume. Antibodies are then added to LNPs and incubated at 4 degrees Celsius overnight to achieve 50 antibodies per liposome as shown in **Supp Fig 1.** When incubation was completed, the liposomes antibody mixture was eluted through a Sepharose column. Binding efficiency of antibodies to liposomes was determined using fluorescence plate reader. Once successful conjugation was determined, the mixture was concentrated using Amicon filters by centrifuging at 3200 x g for 40 minutes. After concentrating the antibody-conjugated LNPs, the particle concentration (number of particles per mL) was measured using nanoparticle tracking analysis (Nanosight), RiboGreen assay, and DLS measurements are performed as mentioned above.

### Animal Studies

#### Animal study protocols

All animal studies were performed in strict accordance with Guide for the Care and Use of Laboratory Animals as adopted by the National Institute of Health and approved by University of Pennsylvania Institutional Animal Care and Use Committee. All animal experiments used male C57BL/6 mice at 6-8 weeks old (Jackson Laboratories). The mice were maintained at 22-26°C and adhered to a 12/12h dark/light cycle with food and water ad libitum.

#### Nebulized LPS Model

Mice were exposed to nebulized LPS in a ‘whole-body’ exposure chamber, with separate compartments for each mouse (MPC-3 AERO; Braintree Scientific, Inc.; Braintree MA). To maintain adequate hydration, mice were injected with 1mL of sterile saline, 37C, intraperitoneally, immediately before exposure to LPS. LPS (L2630-100mg, Sigma Aldrich) was reconstituted in PBS to 10mg/mL and stored at -80C until use. Immediately before nebulization, LPS was thawed and diluted to 5mg/mL with PBS. LPS was aerosolized via mesh nebulizer (Aerogen, Kent Scientific) connected to the exposure chamber (NEB-MED H, Braintree Scientific, Inc.). 5mL of 5mg/mL LPS was used to induce the injury. Nebulization was performed until all liquid was nebulized (∼20 minutes).

#### Biodistribution Measurement

For biodistribution, healthy or nebulized lipopolysaccharide (neb-LPS) treated mice were intravenously injected with Indium-111 radiolabeled aPECAM, DOTAP, or Combined LNPs at a 7.5 ug mRNA dose as previously described^53^. Nanoparticles were produced as described above with 0.1 mol% of 18:0 PE-DTPA (a chelator-containing lipid) using metal free buffers. Trace metals were removed from the buffers using a Chelex 100 resin, per manufacturer’s instructions, to prevent unwanted occupancy of the chelator. In-111 chloride was added to the nanocarriers at a specific activity of 1 uCi of In-111 per 1 umol of lipid. The mixture was incubated at room temperature for 30 minutes. Then, unincorporated In-111 was removed using a Zeba Spin desalting column. The removal of unincorporated In-111 was verified using thin film chromatography (TLC). A 1 uL sample of nanoparticles was applied to the stationary phase (silica gel strip). The strip was placed in the mobile phase of 10mM EDTA until the solvent front was 1 cm from the end of the strip (∼10 minutes). The strip was cut 1 cm above the initial sample location. In-111 chelated to the nanoparticles stays at the origin, while unchelated In-111 travels with the solvent front. The activity in each section was measured using a gamma counter. The percent of In-111 chelated to the nanoparticles was calculated as the activity in the origin strip divided by the total activity in both strips. For all experiments, >95% of In-111 was chelated to the nanoparticles.

Briefly, for neb-LPS treated groups, mice were nebulized four hours prior to injection of nanocarrier as described above. At 10 minutes, 30 minutes or 1 hr after injection with liposomes, mice were euthanized. Blood, lungs, liver, heart, kidney, spleen, injection site and syringe were collected for analysis. Based on a known injected dose of radioactivity, quantified by Wallac 2470 Wizard gamma counter (PerkinElmer Life and Analytical Sciences-Wallac Oy, Turku, Finland), this was used determine the biodistribution by calculating the percent injected dose per gram of tissue.

#### Intra-arterial biodistribution measurement

To perform intra-arterial injections, animals were first anesthetized with ketamine/xylazine, then the right common carotid artery was isolated and cannulated with catheters (Instech) coated with sodium citrate. The common carotid artery bifurcates into the external carotid artery, which supplies blood to the face and neck, and the internal carotid artery, which supplies blood to the brain. To restrict the delivery of the injectate to the head and neck, the external carotid artery was occluded, directing the injectate flow through the internal carotid artery and into the brain vasculature. 100uL of radiolabeled LNPs were infused in around 1 minute.

#### Firefly Luciferase RNA Expression Studies

Luciferase mRNA LNPs fabricated as described above were injected into mice for a circulation time of 4 hours. Select organs were then flash frozen until the day of analysis or homogenized immediately. Samples were suspended in 900 uL of homogenization buffer (5mM EDTA, 10mM EDTA, 1:100 diluted stock protease inhibitor (Sigma), and 1x (PBS), samples were then loaded with a steel bead (Qiagen), then placed in a tissue homogenizer (Powerlyzer 24, Qiagen) using the following settings: Speed (S) 2000 rpm, 2 Cycles (C), T time 45 sec, and pause for 30 sec). After this, 100 uL of lysis buffer (10% Triton-X 100 and PBS) was added into each tube and then allowed to incubate for 1 hr at 4C. After this, they were immediately transferred into fresh tubes, and sonicated, using a point sonicator to remove in excess DNA, using an amplitude of 30%, 5 cycles of 3 secs on/off. After this, samples were then centrifuged at 160,000 x g for 10 minutes. The resultant lysate is either frozen or prepared for luminometry analysis.

For luciferase expression 20 uL of undiluted sample was loaded onto a black 96 well-plate then 100uL luciferin solution (Promega) added immediately before reading on a luminometer (Wallac). Last, a Lowry assay (Bio-Rad) is performed according to manufacturer specification using diluted samples, specifically a 1:40 dilution for lung and spleen tissues and a 1:80 dilution for liver tissues. Final luminescence readings were then normalized based on total protein concentration obtained from Lowry Assay.

#### Cell type distribution and flow cytometry

C57/BL6 mice were injected with a 7.5 ug mRNA dose of fluorescently labeled aPECAM, DOTAP or Combined LNPs and euthanized after 30 minutes (n=3 per group). Lungs were collected and triturated before incubating in a digestive solution of 2 mg/mL collagenase and 100uL of 2.5mg/mL DNase for 45 minutes, to prepare a single cell suspension. After incubation, the cells were strained through a 70uL cell strainer and washed with PBS. The supernatant was discarded and ACK lysis buffer was added for 5 minutes on ice to lyse any remaining red blood cells. After cell lysis, the cells were washed again at 1224 rpm for 5 minutes at 4 degrees Celsius and then counted using an automated cell counter (Countess, Thermo Fisher) to obtain the cell concentrations. The goal was to achieve a concentration of 1 x 10^6^ cells/mL. After cell counting, the suspension was washed with FACS buffer and incubated with Fc block for 15 minutes, followed by 30 minute antibody incubation using CD45, CD64, and Ly6G to identify various leukocytes and CD31 and EPCAM to identify non-leukocyte populations. Fluorophores and markers used for identification of cell types as well as the gating strategy are summarized **in Supp. Fig. 2 and Supp. Table 2**. After antibody incubation was completed, the cells were fixed with 4% PFA prior to analysis on flow cytometer (LSR Fortessa, BD BioSciences).

## Results

### LNPs with physicochemical+affinity targeting achieve markedly improved lung uptake and mRNA expression

To begin these studies, we chose our base LNP formulation to have the ionizable lipid cKK-E12, which others in the field have shown drives high expression from mRNA cargo ^54–56^. We formulated affinity-moiety-targeted LNPs by conjugating aPECAM antibody to the LNP surface (herein referred to as “aPECAM LNPs”). We fabricated LNPs with physicochemical tropism to the lungs by adding in a cationic lipid, DOTAP, as published previously (herein referred to as “DOTAP LNPs”). “Combined LNPs” were formulated by conjugating aPECAM antibody to the surface of DOTAP LNPs (**Fig 1C**). We added a radiotracer lipid (Indium-111-DTPA-PE) to the LNPs and intravenously (IV) injected into healthy mice at a dose of 7.5μg mRNA per mouse. After allowing the LNPs to circulate for 30 minutes, the amount of radioactivity in each organ was measured and recorded as the percent of injected dose per gram of tissue (%ID/g). We found aPECAM and DOTAP LNPs achieved essentially identical lung uptake, despite different targeting mechanisms. LNPs combining the two targeting techniques localize to the lungs >2-fold more than either DOTAP or aPECAM LNPs (**Fig 1A**). Similarly, Combined LNPs have a >2-fold higher lung-to-liver localization ratio than aPECAM or DOTAP LNPs (**Fig 1B**).

To examine protein expression induced by mRNA delivered with each formulation, we fabricated aPECAM, DOTAP, and Combined LNPs loaded with luciferase mRNA. LNPs were IV-injected into healthy mice, and the mice were sacrificed 4 hours later for measurement of luciferase activity in each organ. Combined LNPs have a >2-fold and >1.6-fold higher luciferase expression in the lungs compared to aPECAM or DOTAP LNPs, respectively (**Fig 1D**). Combined LNPs also have a 10-fold and 5-fold higher lung-to-liver luciferase expression ratio than aPECAM and DOTAP LNPs, respectively (**Fig 1E**). Lung-expression-to-uptake ratios were obtained by dividing the normalized luciferase expression values in Fig 1E by the %ID/g values in Fig 1A (**Fig 1F**). We found no significant difference in the expression-to-uptake ratio between formulations, indicating that Combined LNPs’ advantage in luciferase expression is due entirely to their improved lung localization (deposition of nanoparticles in the lung), rather than an advantage in the amount of luciferase expression produced by each LNP that stays in the lungs.

### Cell types targeted by aPECAM, DOTAP, and Combined LNPs

To investigate differences in the cell types that take up our 3 different LNP formulations, we used flow cytometry to trace fluorescent aPECAM, DOTAP, and Combined LNPs in cells isolated from lungs 30 minutes after IV injection of the LNPs (**Fig 2A**, gating strategy shown in **Supp Fig 2)**.

**Figure 2:**
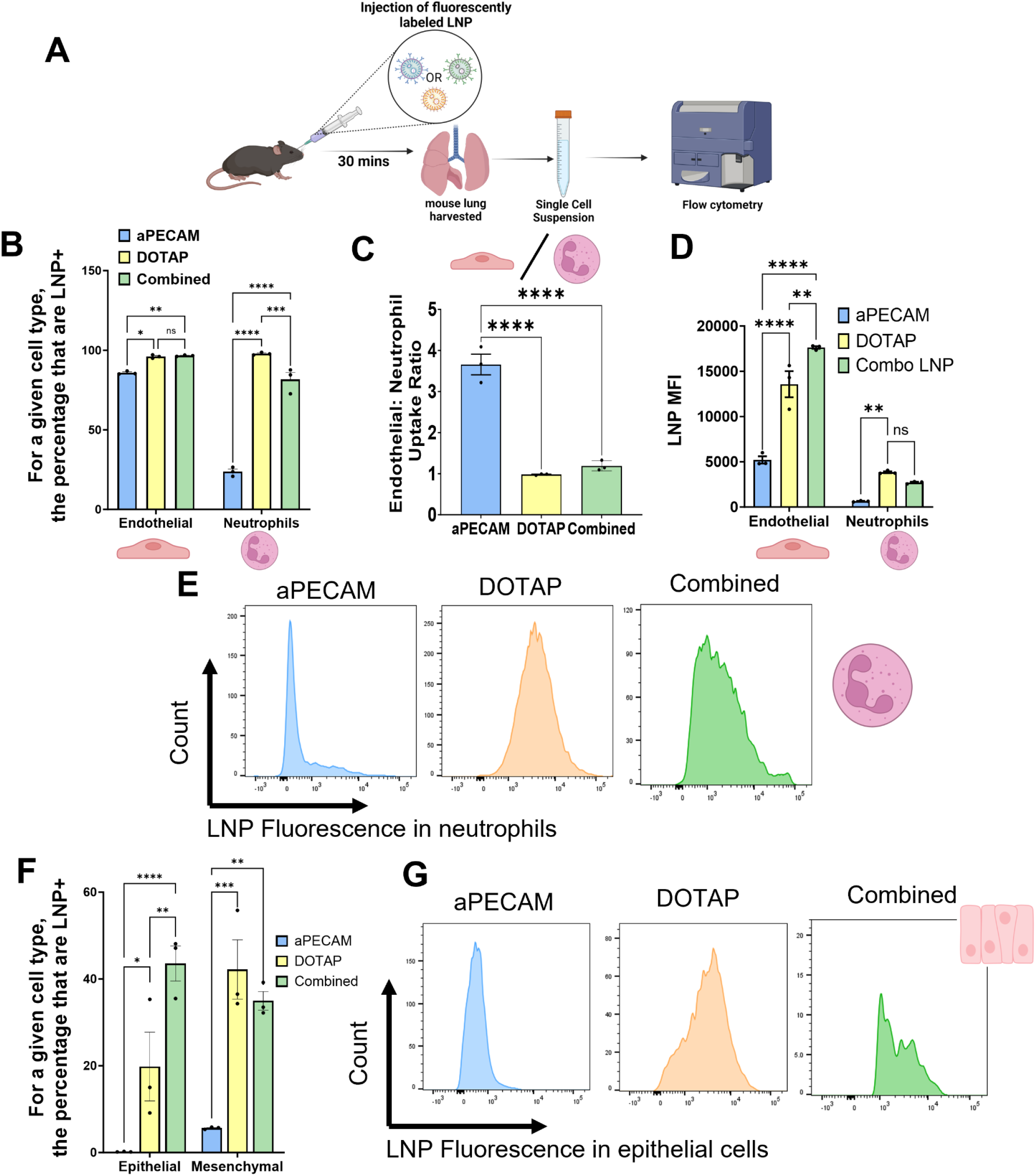
Combined LNPs have enhanced uptake in epithelial and mesenchymal cells *in vivo* in the lung, but also high uptake by neutrophils. **A.** Schematic showing the flow cytometry protocol used, where fluorescently labeled aPECAM, DOTAP, or Combined LNPs were IV injected into mice at a dose of 7.5 ug RNA, allowed to circulate for 30 min, before sacrificed and the lungs prepared as single-cell suspensions. **B.** Gating first by cell type e.g., endothelial cells and neutrophils, we then looked at the percentage of each cell type that was positive for aPECAM, DOTAP, or combined LNPs. Data shows enhanced neutrophil uptake of DOTAP and Combined LNPs **C.** Endothelial-to-neutrophil (E:N) uptake ratio of aPECAM, DOTAP, and Combined LNPs. Values greater than 1 indicate endothelial cell preference compared to neutrophils. aPECAM LNPs are shown to have higher endothelial preference. **D.** LNP mean fluorescence intensity (MFI) for LNP formulations show that while DOTAP and Combined LNPs have a lower E:N ratio compared to aPECAM LNPs, the endothelial cells that take up these LNPs do so highly. E. Representative histograms of LNP fluorescence in neutrophils **F.** Gating first by cell type (as in B), and then plotting LNP positivity among epithelial and mesenchymal cells. These cells are further away from the vascular lumen than endothelial cells and neutrophils, and generally harder to access via intravenous administration. DOTAP and more so Combined LNPs are able to cross and be taken up these two cell types, with Combined LNPs achieving higher epithelial uptake than both DOTAP and aPECAM LNPs. **G.** Representative histograms of epithelial cells showing high LNP fluorescence by DOTAP and Combined LNPs compared to aPECAM LNP. **Statistics:** n=3 and data shown represents mean ± SEM. Comparisons between groups were made using 1-way ANOVA or 2-way ANOVA with Tukey’s post-hoc test where appropriate. *=p<0.05, ***=p<0.001, ****=p<0.0001.

We determined the percentage of endothelial cells and neutrophils that take up aPECAM, DOTAP and Combined LNPs (**Fig 2B**). All three formulations achieve high uptake in endothelial cells (>85% of endothelial cells were LNP-positive), with a small, but significant advantage to DOTAP and Combined LNPs over aPECAM LNPs. Neutrophils took up DOTAP and Combined LNPs in significant quantities (97% and 80% of neutrophils were positive for DOTAP and Combined, respectively), whereas only 23% of neutrophils were positive for aPECAM LNPs. Endothelial-to-neutrophil uptake ratio (**Fig 2C**) shows that aPECAM LNPs have clear endothelial preference compared to DOTAP and Combined LNPs. While aPECAM LNPs had clear endothelial preference over neutrophils, mean fluorescence of DOTAP and Combined LNPs was overall higher in endothelial cells compared to aPECAM LNPs (**Fig 2D**). A large fraction of neutrophils took up DOTAP and Combined LNPs but the MFI data show that these LNPs were taken up in significantly greater concentrations by endothelial cells, compared to neutrophils. This is reflected in the histograms of aPECAM, DOTAP, and Combined LNP fluorescence in neutrophils (**Fig 2E**). In summary, aPECAM LNPs are more endothelial-specific than DOTAP-containing LNPs, with DOTAP-containing LNPs being taken up by a high fraction of neutrophils, but with only a moderate amount of uptake per cell, as compared to endothelial cells.

We similarly assessed monocytes/macrophages, lymphocytes, and dendritic cells (**Supp.** Fig 3A and F). DOTAP and Combined LNPs were taken up in monocytes/macrophages at significantly higher rate than aPECAM LNPs. But uptake in monocytes/macrophages was low, with only 30% of monocytes and macrophages taking up DOTAP and Combined LNPs. Uptake in other leukocytes was similarly low for all LNP formulations.

We assessed LNP uptake in alveolar mesenchymal and epithelial cells, cells that are further away from the alveolar capillary lumen (**Fig 2F**). DOTAP-containing LNPs can reach these deeper cell types^23,24^. 20% and 40% of epithelial cells take up DOTAP and Combined LNPs, respectively. 40% and 35% of mesenchymal cells take up DOTAP and Combined LNPs, respectively. But ∼0% of epithelial cells and about 6% of mesenchymal cells take up aPECAM LNPs. Histograms of epithelial uptake further depict this increase in epithelial delivery (**Fig 2G**). Taken together, we show that while aPECAM LNPs have higher endothelial specificity compared to DOTAP and Combined LNPs, but these latter formulations have the benefit of penetrating to epithelial and mesenchymal cells. There is a synergistic effect between anti-PECAM targeting and cationic lipid tropism on epithelial uptake. However, it is important to note the increased uptake of DOTAP-containing LNPs by neutrophils and monocytes/macrophages, and its implications for the inflammatory and thrombotic responses to LNPs.

### Effects of pathology on organ and cell type targeting

We used the most common model for ARDS, nebulized LPS (neb-LPS)^57,58^, to investigate how acute lung inflammation affects LNP lung targeting and cell type specificity. Radiolabeled or fluorescent LNPs were administered IV 4 hours after neb-LPS, at which point inflammatory processes and neutrophilic infiltration into the lung had developed (**Fig 3A**).

**Figure 3:**
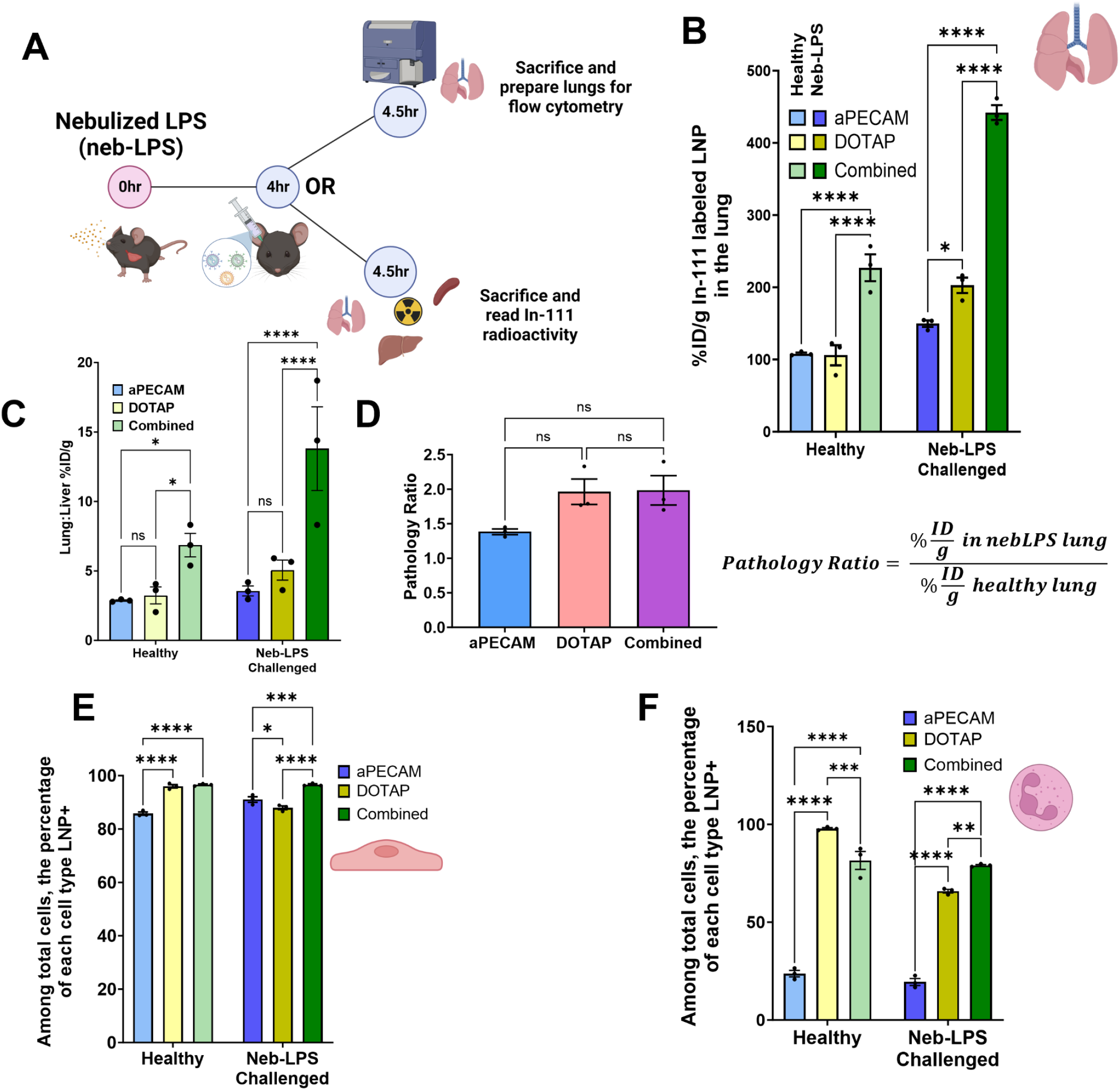
Combined LNPs show superior lung localization in a mouse model of acute lung inflammation. **A.** Schematic of nebulized-LPS injury and resulting assays performed, briefly 4 hours after neb-LPS injury, mice are either injected with fluorescently labeled or Indium-111 (In-111) radiolabeled nanoparticles, allowed to circulate for 30 minutes before sacrifice to determine either cell type distribution or whole organ particle localization. **B.** Comparative biodistribution between healthy and neb-LPS challenged mice given aPECAM, DOTAP, and combined LNPs show enhancement of localization for all formulations to the lung following neb-LPS challenge. Combined LNPs localize to the lungs 3-fold and 2.25-fold more than aPECAM and DOTAP LNPs respectively following neb-LPS challenge. **C.** Comparison of lung-to-liver ratios between healthy and neb-LPS challenged mice, shows significant enhancement to lung localization after neb-LPS challenge for all formulations but especially for the combined formulation, getting an almost 2-fold improvement in lung targeting. Similarly, combined LNPs have a >2-fold higher lung to liver ratio compared to aPECAM or DOTAP LNPs also after neb-LPS injury. **D.** Quantification of the effect of pathology lung uptake and delivery aPECAM, DOTAP, and Combined LNP formulations. Dividing the %ID/g of LNP in the lung after neb-LPS by the %ID/g in healthy lung shows that aPECAM, DOTAP, and Combined LNPs increase localization between 1.5-2 fold increase in lung localization. **E.** Gating first by cell type (endothelial cells) then identifying the percentage that are LNP positive, shows that pathology does not influence uptake of LNPs by endothelial cells. **F.** Similarly, this is the case for neutrophils. **Statistics:** n=3 and data shown represents mean ± SEM. Comparisons between groups were made using 1-way ANOVA or 2-way ANOVA with Tukey’s post-hoc test where appropriate. *=p<0.05, ***=p<0.001, ****=p<0.0001.

In radiotracer data, Combined LNPs had a 2-fold improvement in lung uptake in neb-LPS mice vs. healthy animals (**Fig 3B** and **Supp Fig 1**). Lung uptake of Combined LNPs in neb-LPS mice was also 2-fold higher than that of aPECAM or DOTAP LNPs. Lung-to-liver uptake ratio follows a similar trend (**Fig 3C**), showing high lung specificity of Combined LNPs after neb-LPS injury. We define a “pathology ratio” as lung uptake in neb-LPS mice divided by lung uptake in healthy mice. aPECAM, DOTAP, and Combined LNPs had statistically similar pathology ratios, with neb-LPS causing a ∼1.5 to 2-fold increase in lung uptake (**Fig 3D**).

Despite increased lung uptake in radiotracer data, in flow cytometry data, there was no increase in either endothelial cells or neutrophils positive for LNPs when comparing neb-LPS to healthy mice (**Figs 3E-F**).

Epithelial and mesenchymal uptake of DOTAP and Combined LNPs was similarly conserved (**Supp Fig 3G**). There was an increase in monocyte and macrophage uptake of DOTAP and Combined LNPs, from 30% positive in healthy mice to 40% in neb-LPS (**Supp Fig 3B and F**). However, mean LNP fluorescence for these cell types do not show increased LNP uptake, indicating that they are not the source for the increase in lung uptake (**Supp Fig 3D**).

### DOTAP and Combined LNPs provide unprecedented delivery to the brain via intra-arterial injection

Since the lungs possess the first capillary bed downstream of an IV injection, we used intra-arterial (IA) injection in the right carotid artery to see if a first pass effect may; a) have contributed to lung uptake of our LNPs; b) allow for increased LNP uptake in the brain after intracarotid injection (**Fig 4A**). After IA injection, radiolabeled aPECAM, DOTAP, and Combined LNPs were allowed to circulate for 30 min before assessing biodistributions.

**Figure 4:**
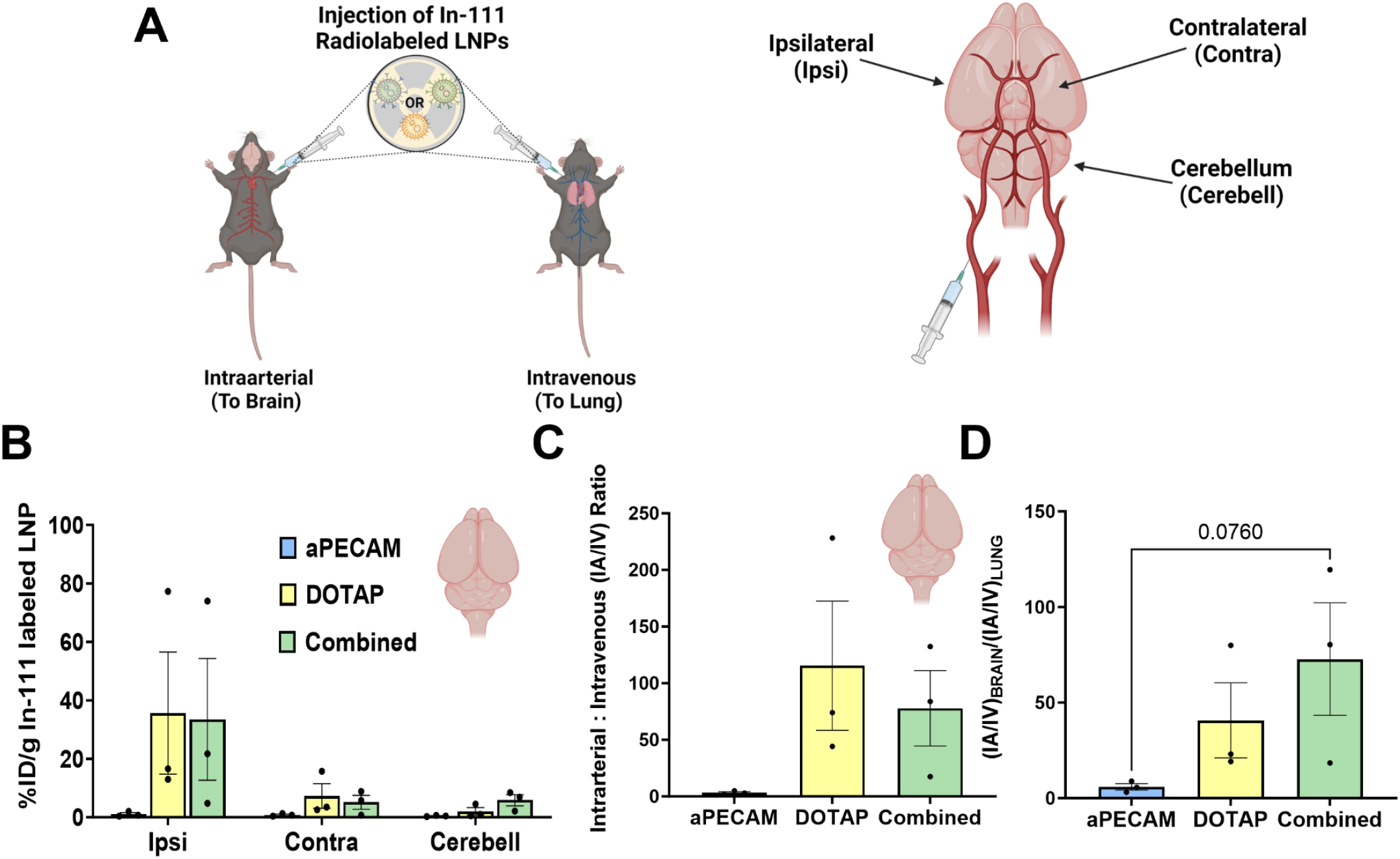
DOTAP and Combined LNP formulations are dependent on first pass privilege for brain localization, but provide unprecedented targeting to the brain. A test for targeted therapeutics specificity to the lung is to measure the dependence of first pass effect as the lung is the first major capillary bed targeted therapeutics face. **A.** To eliminate first pass effect, In-111 radiolabeled aPECAM, DOTAP, and Combined LNPs were injected intra-arterially (IA) into the right carotid artery and allowed to circulate for 30 minutes, the animals were then sacrificed and organs measured for radioactivity, this was compared to direct intravenous delivery (IV) via retro orbital injection. Segments of the brain were dissected as shown in **A (right)** identifying ipsilateral (same side of injection), contralateral (opposing side of injection), and the cerebellum. **B.** Following IA injection of aPECAM, DOTAP, and Combined LNPs we show enhanced localization by DOTAP and Combined LNPs in the brain, specifically in the ipsilateral cerebrum (same side where the LNP was injected). **C.** Directly comparing the %ID/g in the brain after IA injection to %ID/g after IV injection highlights that for brain localization, where IA administration gives the brain capillary bed first pass privilege, enhances delivery by DOTAP and Combined LNPs by an order of magnitude. **D.** We next compare this IA:IV ratio in the brain to that of the lung to better evaluate intraarterial first pass effect dependence on targeting to the brain. **Statistics.** n=3 and data shown represents mean ± SEM. Comparisons between groups were made using 1-way ANOVA, where appropriate with Tukey’s post-hoc test.

DOTAP LNPs have increased uptake in the lungs after IA injection, compared to IV injection (**Supp Fig 4A and B**). This may be because IA injection provides the DOTAP LNPs an extra cardiac cycle before lung exposure, allowing for accumulation of additional opsonins (see Discussion for further hypotheses). aPECAM and Combined LNPs’ lung uptake was unaffected by the injection method. Our results indicate a first-pass effect is not the main reason for lung uptake of our targeted LNPs.

After IA injection in the right carotid artery, we observed high uptake of DOTAP and Combined LNPs in the brain, but no such uptake with aPECAM LNPs. The uptake of DOTAP and Combined LNPs in the right (ipsilateral to injection site) hemisphere was 35% ID/g, which is, to our knowledge, the highest reported delivery to the brain (**Fig 4B**)^53,59^. IA injection also achieves a moderate increase in DOTAP and Combined LNP uptake in the contralateral side and the cerebellum. Comparing ipsilateral brain uptake of LNPs after IA vs. IV delivery shows IA delivery increases brain delivery ∼50-100-fold (**Fig 4C**). This implies that DOTAP-containing LNP brain uptake is almost entirely driven by a first-pass effect. Comparing the IA-to-IV ratio for brain uptake vs lung uptake shows a greater first-pass effect for brain delivery than for lung delivery of DOTAP-containing LNPs, especially Combined (**Fig 4D**).

## Discussion

Two classes of targeting methods have arisen to achieve LNP delivery to specific cell types and organs: affinity moiety targeting ^29,31,32,35^ and physicochemical tropism ^19,23–25,60^. In this study, we directly compared these two methods. We largely focused on targeted delivery to the lungs, using aPECAM LNPs to represent affinity moiety targeting, and DOTAP LNPs to represent for physicochemical tropism. We also incorporated aPECAM and DOTAP in the same LNPs, to achieve a “Combined” targeting LNP.

Focusing on the comparison between affinity moiety targeting and physicochemical tropism, the two different targeting strategies produced nearly identical overall lung uptake. We have previously observed that different targeting strategies yield similar lung uptake, in studies with nanoparticles targeted via aPECAM, aICAM^61^, with the technique of RBC-hitchhiking ^62–64^, or with physicochemical tropism to marginated neutrophils^65^. This suggests a saturation value for lung uptake with most kinds of targeting focused on the alveolar capillaries. Combined LNPs overcome this saturation, as discussed below.

Comparing cell type selectivity of affinity moiety targeting vs. physicochemical tropism, aPECAM LNPs have better endothelial specificity than DOTAP LNPs. Endothelial cells are traditionally the cell of interest in lung targeting. However, DOTAP provides delivery to cells outside of the vasculature, such as epithelial and mesenchymal, and thus promises RNA delivery to lung cell types not usually accessed with IV agents.

Most notably, combining aPECAM and DOTAP provides a synergistic benefit to lung uptake. Combined LNPs had > 2x more lung uptake than either aPECAM or DOTAP alone and provided unprecedented delivery to alveolar epithelial cells. And combined LNPs had 2x-higher delivery to inflamed lungs than to healthy lungs. These advantages strongly recommend further investigation of Combined LNPs and their mechanisms of higher uptake, especially in the setting of pathology, such as in pulmonary inflammation.

DOTAP and Combined LNPs strongly localize to the brain following intra-arterial injection. They achieve ∼35% ID/g on the side of the brain ipsilateral to injection site. This level of brain uptake greatly exceeds standard brain targeting methods, including transferrin and VCAM targeting ^53,59^. There was also an increase in uptake in the contralateral brain and cerebellum, but most brain uptake was on the ipsilateral side. Lateral selectivity in brain uptake can be useful for brain injuries that present unilaterally, including stroke and traumatic brain injuries.

Importantly, safety and side effects are critical issues which must be investigated in future studies. As with any developing technology, Combined LNPs will certainly have side effects, and it is important to investigate these and attempt to engineer solutions. Of greatest concern with DOTAP-containing LNPs is that we recently showed that IV LNPs formulated with any of various cationic lipids cause thrombosis in the lungs^66^. It is reasonable to hypothesize that Combined LNPs may cause similar effects. However, our recent work also demonstrated techniques to mitigate clotting induced by cationic lipid LNPs, including anticoagulation and reduction of LNP size. These techniques can and should be applied to Combined LNPs in future studies. It remains to be seen whether these or other engineered solutions will be sufficient to make the risk-benefit ratio meet the criterion for further preclinical development of what appears to be an promising targeting technology.

Additional safety concerns of note include LNP-associated inflammation. We have previously shown that LNPs as a class worsen pre-existing inflammation^67^. Here we show that DOTAP-containing LNPs have greater uptake by innate immune cells, which may lead to even greater inflammation than other LNPs. Our flow cytometry data shows that DOTAP and Combined LNPs are avidly taken up by neutrophils, monocytes, and macrophages, especially in the context of acute lung injury (**Figs 2B, 3F, and Supp Fig 3A, B, and F**). Increased innate immune uptake of LNPs increases the potential for activation of inflammatory pathways and opens the door for later adaptive immunity against the LNPs. Like with other possible toxicities, future studies should aim to engineer solutions to uptake of DOTAP-containing LNPs by innate immune cells, such as adding on “don’t eat me” signals such as CD47-mimetic peptides^68^. On the flip side, DOTAP-LNPs’ uptake by innate immune cells also present new targets for RNA delivery and disease-specific treatments ^58,69–71^.

In summary, this study shows that aPECAM affinity moiety targeting and cationic lipid physicochemical targeting achieve similar levels of IV lung uptake, but target different cell types in the lungs. LNPs combining these two targeting methods (“Combined LNPs”) provide 2x better lung uptake and increased uptake in extravascular cells. Combined LNPs achieve excellent brain uptake when delivered intra-arterially. These findings suggest that combining the two classes of nanoparticle targeting can provide benefits not seen with either targeting method alone.

## Notes

### Competing Interest Statement

The authors have declared no competing interest.

