## SupplementalFigures for "A combination of physicochemical tropism and affinity moiety targeting of lipid nanoparticles enhances organ targeting"

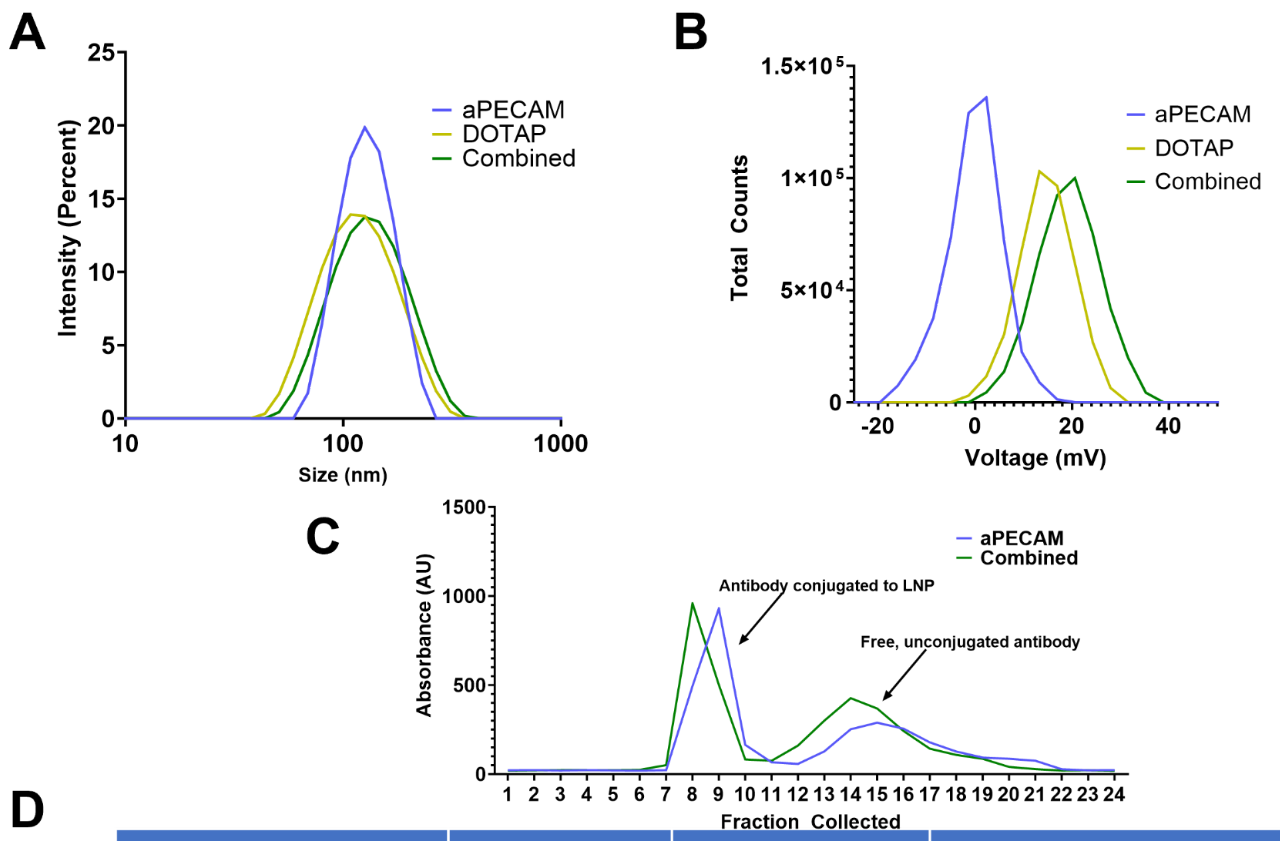


D

| LNP Formulation | Size (nm) | Charge (mV) | Conjugation Efficiency |
| --- | --- | --- | --- |
| aPECAM | 117.8±7.01 | -2.93±0.41 | 57% |
| DOTAP | 114.9±7.47 | 12.37±4.2 | N/A |
| Combined | 123.5±13.9 | 15.56±3.10 | 50% |

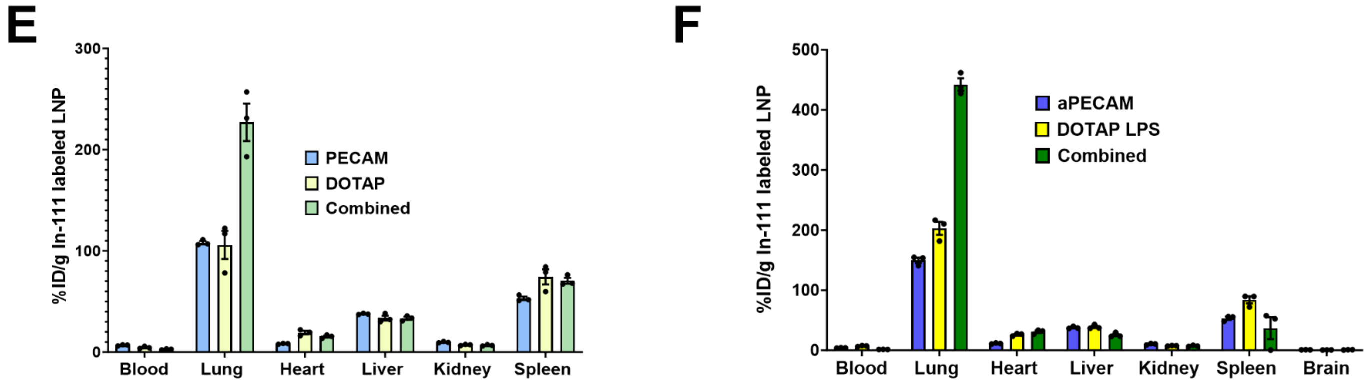


**Supp. Figure 1: Characterization of aPECAM, DOTAP, and Combined LNPs A.** Representative DLS measurements of aPECAM, DOTAP, and Combined LNPs depicting size and **B.** Zeta potential. **C.** Representative graph of elution profile of AF 647 labeled PECAM antibody generated by size exclusion chromatography (SEC) via Sepharose column for aPECAM and Combined LNPs. Due to the larger size of LNP, aPECAM and DOTAP LNPs leave the Sepharose column at fractions 7-10. This is later confirmed by DLS. **D.** Table summarizing size, charge, and conjugation efficiency. To achieve the desired antibody number on the surface of the LNP, this value is taken into consideration and excess antibody is provided. **E.** Biodistribution for aPECAM, DOTAP, and Combined LNPs highlighting all major clearance organs. **F.** Effect of pathology on overall organ localization of aPECAM, DOTAP, and Combined LNPs.

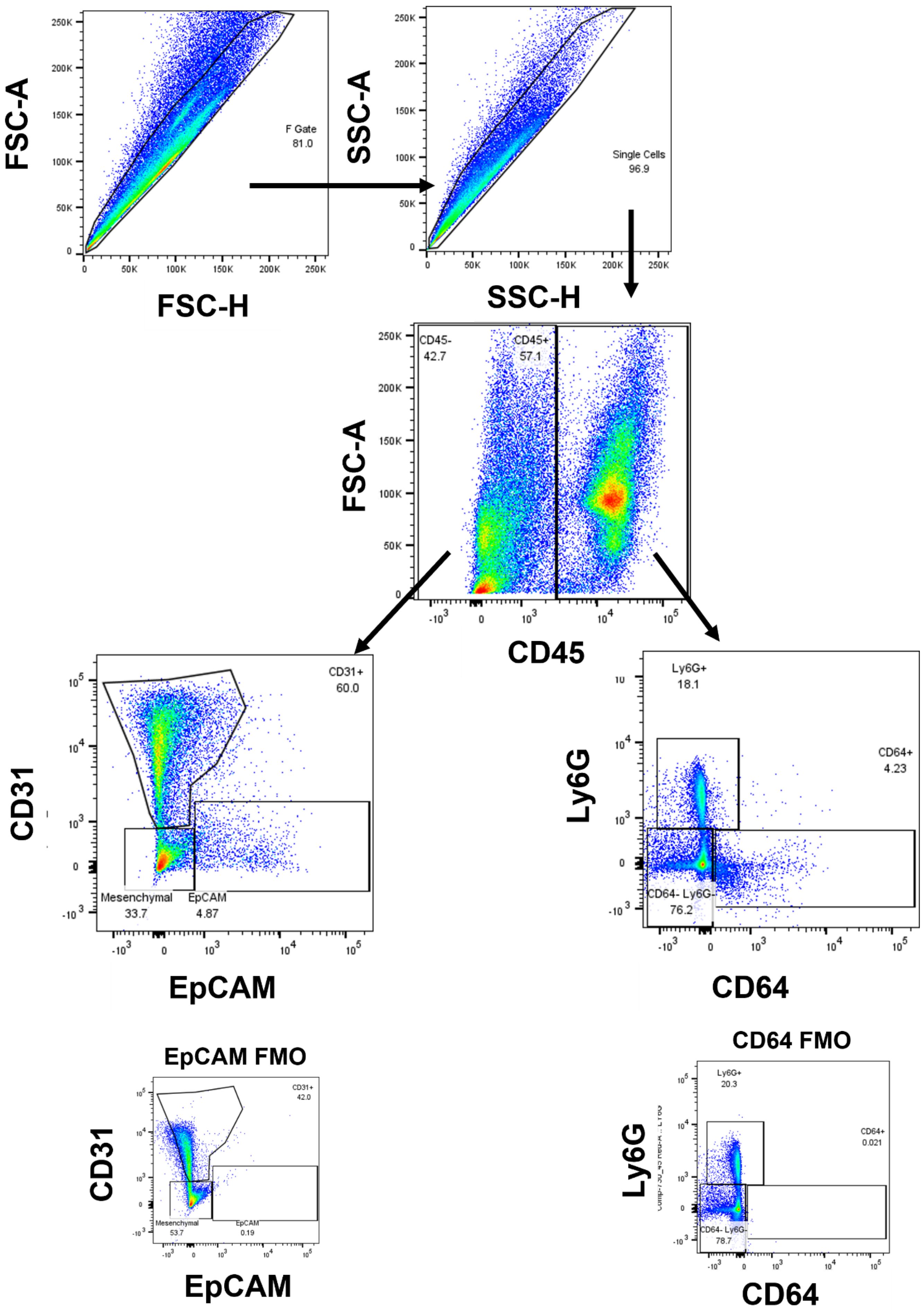
FSC-A
FSC-H
F Gate
81.0
SSC-A
SSC-H
Single Cells
96.9
FSC-A
CD45
CD45-
42.7
CD45+
57.1
CD31
EpCAM
CD31+
60.0
Mesenchymal
33.7
EpCAM
4.87
Ly6G
CD64
Ly6G+
18.1
CD64+
4.23
CD64- Ly6G-
76.2
EpCAM FMO
CD31
EpCAM
CD31+
42.0
Mesenchymal
53.7
EpCAM
0.19
CD64 FMO
Ly6G
CD64
Ly6G+
20.3
CD64+
0.021
CD64- Ly6G-
78.7

**Supp Figure 2: Gating Strategy to determine cell types that take up aPECAM, DOTAP, and Combined LNPs.** Gating strategy used to determine specific cell types assessed in this study. First sorting by Forward scatter, then by side scatter to remove doublets and debris from the analysis. We then sort by CD45 negative and positive to identify leukocytes. In the leukocyte arm, we use Ly6G to identify neutrophils and CD64 to identify monocytes and macrophages. For non-leukocytes, we use CD31 to identify endothelial cells and EpCAM to identify epithelial cells. For epithelial cells as well as monocytes and macrophages, fluorescence minus one (FMO) controls were generated to positively identify these cell types.

| Lipid | aPECAM (Radiolabeled) | aPECAM (Fluorescent) | DOTAP (Radiolabeled) | DOTAP (Fluorescent) | Combined (Radiolabeled) | Combined (Fluorescent) |
|---|---|---|---|---|---|---|
| cKK-E12 | 50.0% | 50.0% | 25.0% | 25.0% | 25.0% | 25.0% |
| Cholesterol | 38.4% | 38.2% | 18.4% | 18.20% | 18.4% | 18.20% |
| DOPE | 10.0% | 10.0% | 5.0% | 5.0% | 5.0% | 5.0% |
| DSPE-PEG-2k azide | 1.5% | 1.5% | N/A | N/A | .75% | .75% |
| DMG-PEG-2k | N/A | N/A | 1.5% | 1.5% | .75% | .75% |
| DOTAP | N/A | N/A | 50% | 50% | 50% | 50% |
| 18:1 PE TopFluor AF 594 | N/A | 0.3% | N/A | 0.3% | N/A | 0.3% |
| 18:0 PE-DTPA | 0.1% | N/A | 0.1% | N/A | 0.1% | N/A |

**Supp Table 1**: Molar percentages used in fabrication of both radiolabeled and fluorescently labeled aPECAM, DOTAP, and Combined LNP formulations N/A signifies 0% of that lipid was added to that formulation.

| Marker | Cell Type | Color | LSR Fortessa Detector |
|---|---|---|---|
| CD45 | General Leukocyte | BUV 395 | UV 380/30 |
| Ly6G | Neutrophil | AF700 | RED 730/45 |
| CD31 | Endothelial | APC | RED 670/14 |
| EpCAM | Epithelial | BV 711 | VIOLET 710/50 |
| CD64 | Monocyte/ Macrophage | PE-Cy7 | YG 780/60 |
| Fluorescent Lipid | LNP Formulations | AF 594 | YG 610/20 |

**Supp Table 2: Flow cytometry markers and laser detectors:** Surface markers used for flow cytometry identification. Provided here are the fluorophores used as well as the detectors used on the LSR Fortessa for identification.

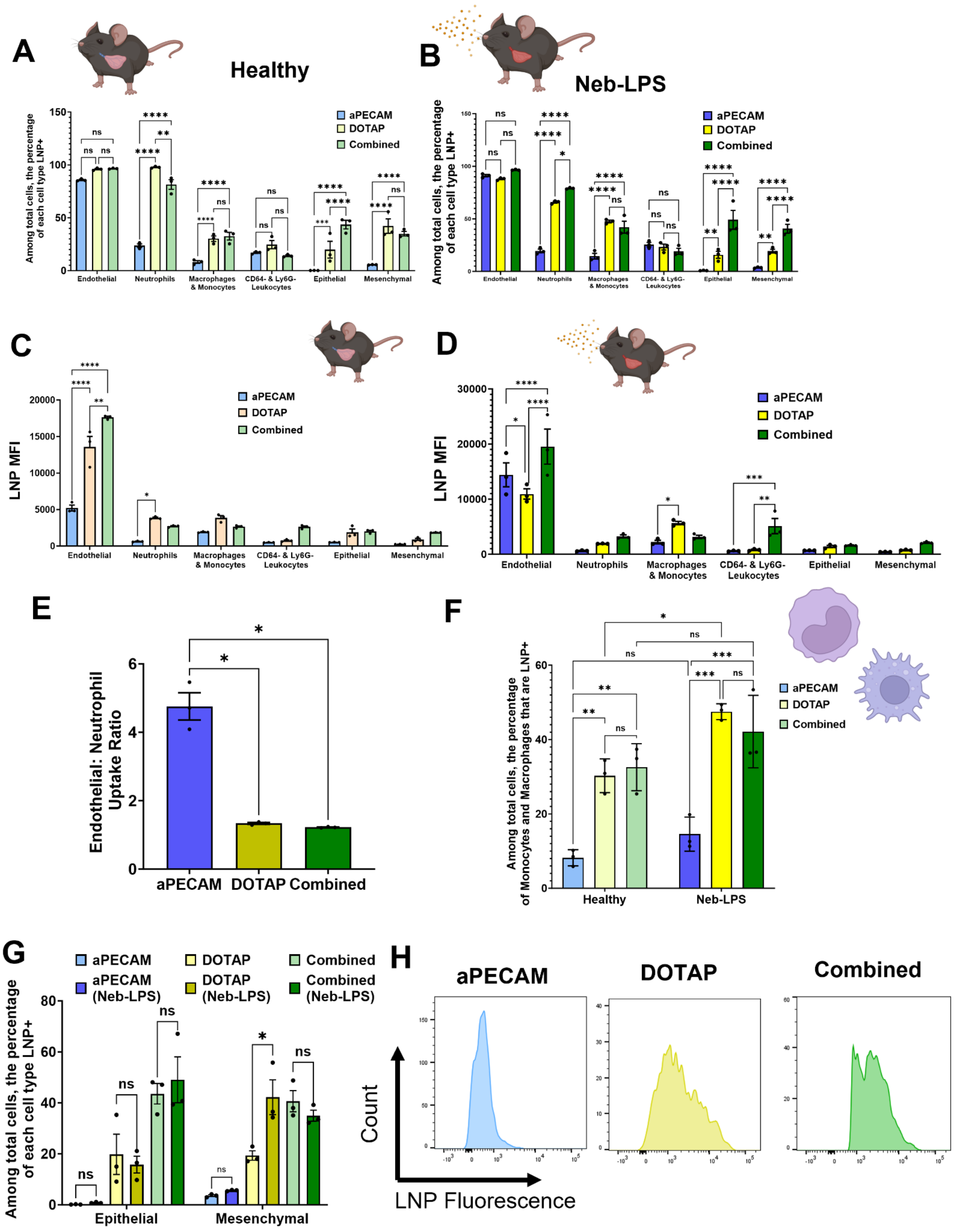


**Supp. Fig 3: Effects of pathology on cell specific uptake A.** First by gating among total cells, then assessing the percentage of those cells that are LNP positive. Here we show all of the cell types that were analyzed, and similarly for **B** after neb-LPS injury. **C.** LNP MFI shows that while there was no significant difference in endothelial cell uptake in healthy animals as shown in **A**. However, MFI results show a higher fluorescence intensity for DOTAP and Combined LNPs showing that endothelial cells have a higher preference for these formulations. **D.** Interestingly, only Combined LNPs maintain this heightened endothelial preference after neb-LPS challenge. It is also worth noting that Combined LNPs also elicit increased uptake by leukocytes. **E.** After the neb-LPS challenge, aPECAM LNPs still maintain their endothelial specificity compared to DOTAP and Combined LNPs. **F.** Comparing healthy to neb-LPS challenged mice and then assessing by flow cytometry the percentage of monocytes and macrophages that are LNP positive. Interestingly, compared to aPECAM LNPs, both DOTAP and Combined LNPs are more highly taken up by monocytes and macrophages in both healthy and neb-LPS challenged mice. Additionally, only DOTAP LNPs show an increase in monocyte and macrophage uptake from healthy to neb-LPS challenged mice, likely from increased immune activation from the injury. **Statistics**. n=3 and data shown represents mean ± SEM. Comparisons between groups were made using 1-way ANOVA or 2-way ANOVA with Tukey's post-hoc test where appropriate. *=p<0.05,**=p<.01 ,***=p<0.001, ****=p<0.0001.

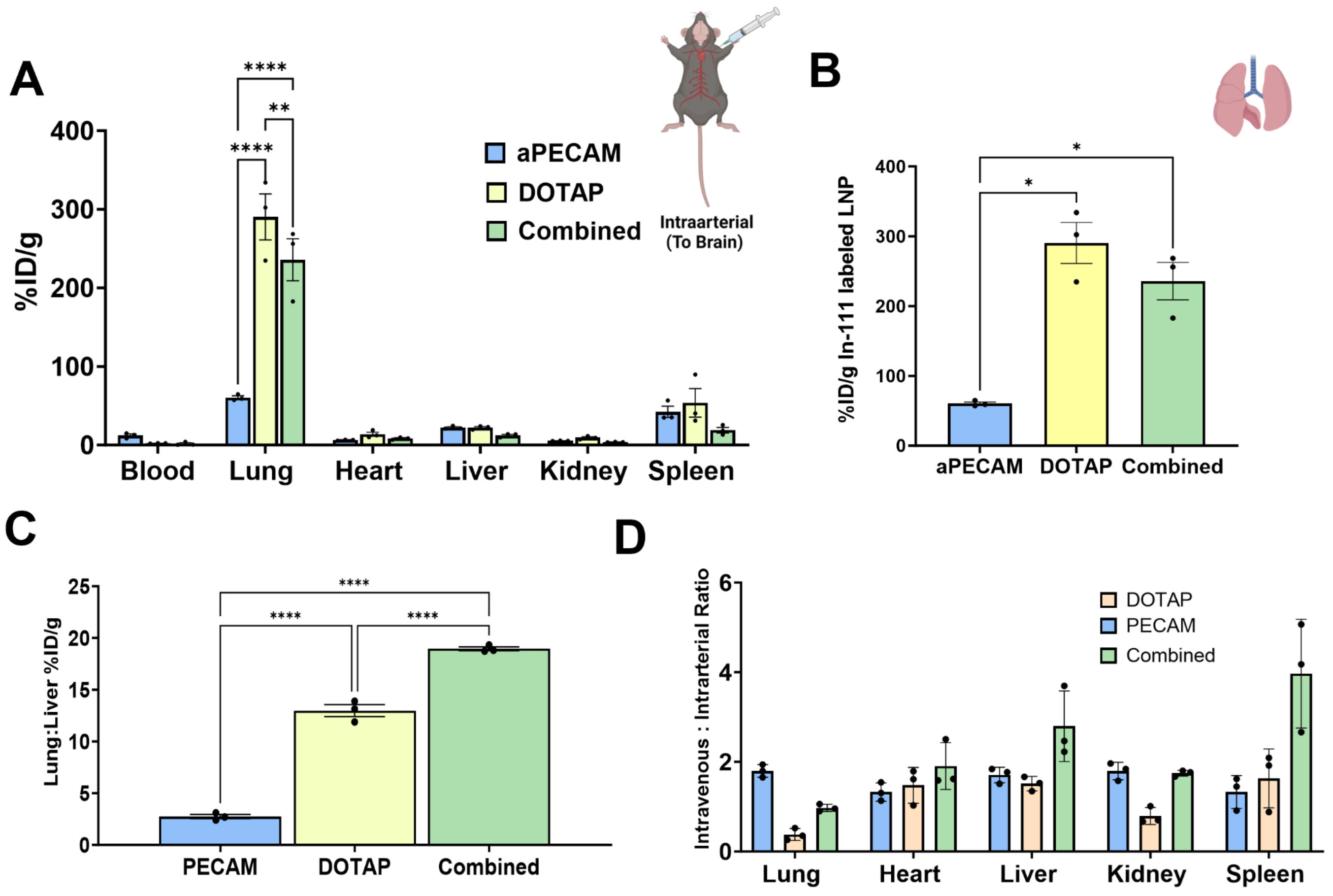


**Supp. Fig 4: Effect of intra-arterial versus intravenous injection of aPECAM, DOTAP, and Combined LNPs on Lung Targeting. A.** Intra-arterial injection of aPECAM, DOTAP, and Combined LNPs for all other organs. We see that compared to aPECAM LNPs, DOTAP and Combined LNPs show even higher specificity toward the lung. This is highlighted in **B**. **C.** This is further depicted by the lung-to-liver ratio which shows that Combined LNPs have enhanced specificity relative to the liver and outcompetes both aPECAM and DOTAP LNPs. **D.** Here, we compare the effect of intravenous injection to intra-arterial injection. A values >1 indicate preference for intravenous delivery, while values <1 show preference for intra-arterial delivery for delivery to that organ. We see here that for lung delivery, aPECAM LNPs prefer intravenous delivery, suggesting these LNPs are influenced by first pass effect. Further DOTAP and Combined LNPs were found to not be first pass dependent. Interestingly, lung localization for Combined LNPs seems to be agnostic to either mode of injection, however lung localization for DOTAP LNPs seems to be driven by intra-arterial injection, likely from increased circulation time allowing for more protein corona formation. **Statistics**. n=3 and data shown represents mean ± SEM. Comparisons between groups were made using 1-way ANOVA or 2-way ANOVA with Tukey's post-hoc test where appropriate. *=p<0.05,**=p<.01 ,***=p<0.001, ****=p<0.0001.

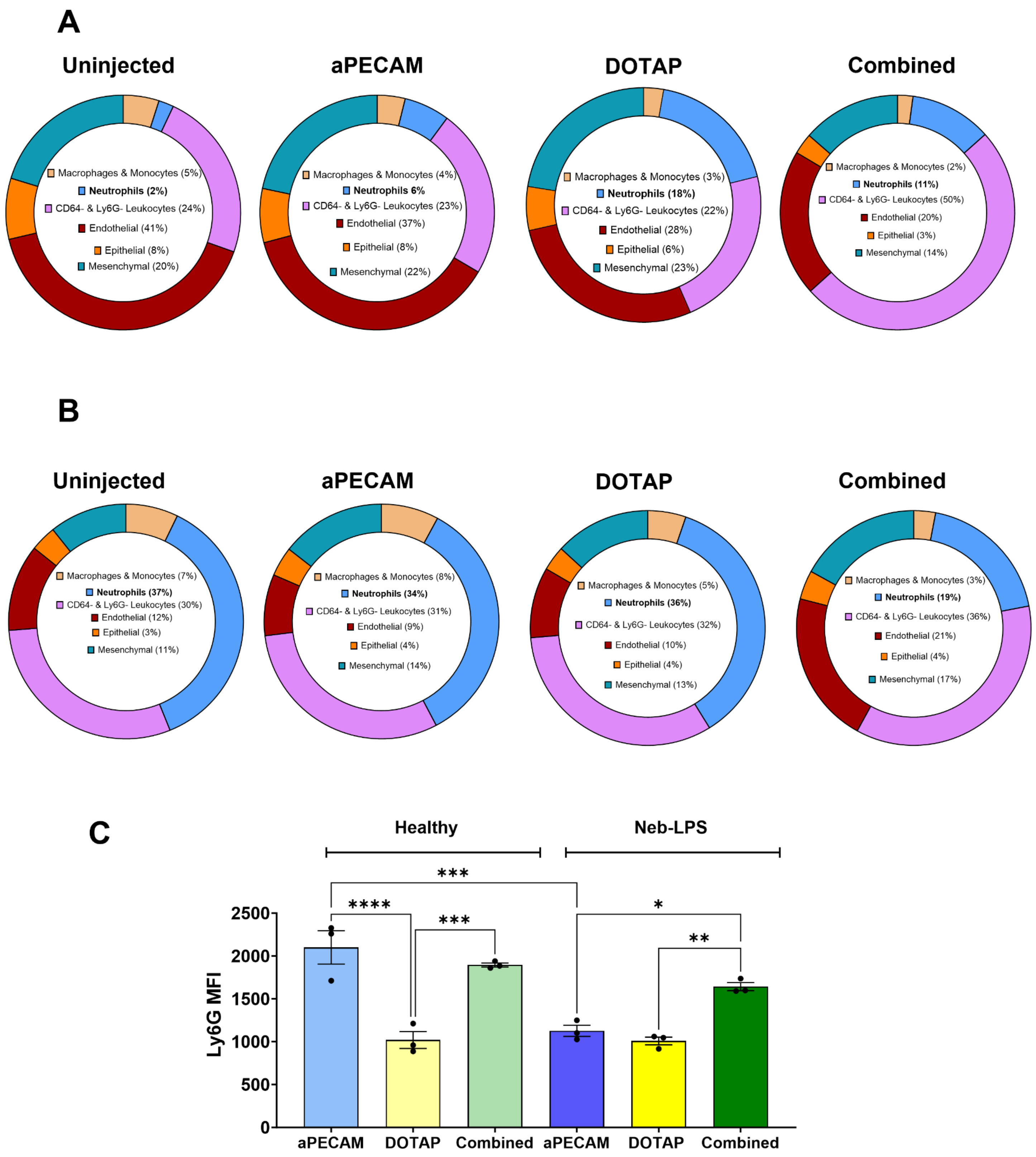


**Supp Fig 5. Effects of LNP formulation on cell type distribution. A.** Comparative cell type distribution of healthy mice after administration of no particles, aPECAM, DOTAP, and Combined LNPs. Data shows that all three formulations significantly increase neutrophil presence in the lung comparative to untreated control, but DOTAP LNP formulation shows the highest increase in neutrophil presence at 18% of total recovered single cells. **B.** After 4 hour

Neb-LPS injury followed by administration of either aPECAM, DOTAP, or Combined LNP formulations, we observe for all formulations and control an increase in neutrophil presence in the lung, as expected for acute lung injury. However, the increase is not as large for Combined LNPs and is markedly lower comparative to control mice administered neb-LPS only suggesting possible clearance of neutrophils from the lung. **C.** Comparison of Ly6G MFI 30 min after administration of either aPECAM, DOTAP, or Combined LNPs. Data shows a significant reduction in Ly6G MFI. This reduction in MFI is suggestive of a left-shift and increase in the presence of immature neutrophils. **Statistics**. n=3 and data shown represents mean ± SEM. Comparisons between groups were made using 1-way ANOVA with Tukey's post-hoc test where appropriate. *=p<0.05,**=p<.01 ,***=p<0.001, ****=p<0.0001.
